# Host Genome Drives the Diversity, Richness, and Beneficial Microbes in the Shrimp Microbiota: A Hologenome Perspective

**DOI:** 10.1101/2024.10.01.616099

**Authors:** Fernanda Cornejo-Granados, Luigui Gallardo-Becerra, Sandra Romero-Hidalgo, Alonso A. Lopez-Zavala, Rogerio R. Sotelo-Mundo, Andres Cota-Huizar, Adrian Ochoa-Leyva

**Affiliations:** Departamento de Microbiología Molecular, Instituto de Biotecnología (IBT), Universidad Nacional Autónoma de México (UNAM) Av. Universidad #2001, Col. Chamilpa, Cuernavaca, Morelos 62210 México; Departamento de Genómica Computacional, Instituto Nacional de Medicina Genómica, Secretaría de Salud (INMEGEN), Periférico Sur No. 4809, México, DF 14610, México; Departamento de Ciencias Químico Biológicas. Universidad de Sonora (UNISON). Blvd., Rosales y Luis Encinas. Hermosillo Sonora 83000 México; Laboratorio de Estructura Biomolecular, Centro de Investigación en Alimentación y Desarrollo, A.C. (CIAD), Carretera Gustavo Enrique Astiazarán Rosas num. 46. Col. La Victoria,, Hermosillo, Sonora, 83304, México; Camarones El Renacimiento SPR de RI. Justino Rubio No. 26, Col Ejidal, Higuera de Zaragoza, Sinaloa 81330 México

## Abstract

Despite the growing understanding of the factors influencing the shrimp microbiome, the impact of host genetics on modulating the intestinal microbiota still needs to be studied. Moreover, the existing studies were typically conducted using animal models and under laboratory conditions. Here, we investigated the effect of two shrimp genetics, fast-growth (Gen1) and disease-resistant (Gen2), on shaping the microbiota of the hepatopancreas and the intestine in open-pond aquaculture farming conditions. First, we identified significant genomic differences between the two genetic lines using genome-wide SNP analysis. Then, the microbiota analysis confirmed that organs had a more substantial impact, explaining 33.9% of the variability, followed by the genetic line, which explained 17.3%. The microbiota of the hepatopancreas was more significantly affected by the genetic line than the intestine. Gen1 exhibited higher richness, diversity, niche breadth, and abundance of beneficial microbes than Gen2, suggesting that Gen1 had a more generalist microbiota. By comparing the microbiota of another set of healthy and diseased shrimps, we confirmed that a higher presence of beneficial microbes was linked to a healthy shrimp status. Additionally, we genotyped and determined the microbiota of wild-type shrimps, proving that they differ from those observed in both genetic lines. Interestingly, ponds with Gen1 had better productivity than Gen2, suggesting a potential link between higher productivity and the microbiota selected by Gen1. Our study highlights the importance of the holobiome perspective in breeding and management programs. It suggests that a specific genetic line and its associated microbiota could be used to select the larvae to be cultivated, improving shrimp aquaculture production.

## Introduction

The influence of the host genome on the microbiome is typically studied using model systems or in laboratory settings. Understanding the interactions between the initial seed microbiota present at birth and the changes during development, diet, and rearing is crucial for gaining insight into diseases and metabolic disturbances (1). The study of microbiota in aquaculture and fisheries is vital for global food production and food security, in addition to human health and animal rearing. According to the FAO, 89% of the aquatic animal production in 2020 was directly used for human consumption (2), and this number is expected to increase due to the growing human population (3).

Crustaceans are the second leading aquaculture product worldwide, with the whiteleg shrimp (*Litopenaeus vannamei*) being the most valuable species. However, shrimp farming faces significant challenges due to diseases, resulting in an estimated annual production loss of 1-4 billion dollars. The increasing demand for shrimp has led the aquaculture industry to explore strategies to enhance production and maintain shrimp health and growth. These strategies include intensifying rearing systemtonss, improving feed formulations, using dietary supplements, or employing microbial consortia to improve water quality. However, these strategies could disrupt the surrounding microbial balance, impacting shrimp health and growth.

The relationship between shrimp health and its microbiota has been extensively studied over the past decade, with mounting evidence indicating a close connection (4–7). Effective management of the shrimp’s microbiota, the surrounding sediments and water reservoirs, is crucial for achieving a sustainable source of high-quality proteins (3) with a lower carbon footprint compared to terrestrial livestock (8). The composition of microbial communities is influenced by various factors, including diet, pre- and probiotics, antibiotics (5), environmental conditions such as farmed or wild-type settings (9), water salinity, and the shrimp’s developmental stage (10).

Currently, agriculture (11), livestock (12), and aquaculture (13) industries are exploring the use of probiotics to establish a healthy microbiota that improves intestinal health, enhances the immune system, and promotes the growth of plants and animals for consumption. However, there are challenges related to the effectiveness of potential probiotics. These bacteria must survive in sufficient numbers, adhere to the host’s intestinal mucosa, withstand environmental conditions, coexist with the existing microbiota in their host, and multiply (13). These factors underscore the host’s role in shaping the microbiota (9,14–17) and how probiotics effective for one animal species may not be optimal for others. This highlights the need for further research in this area.

The concept of a unique microbiota composition for every species, as supported by numerous studies (14,18,19), gained a significant boost in 2008 with the introduction of the hologenome concept by Zilber-Rosenberg and Rosenberg (20). This concept, which defines the total genetic information of the host and its microbiota, has profound implications for the evolution and adaptation of higher organisms. It suggests that the co-evolution of their microbial symbiotic genomes is a key factor. This means that the microorganisms residing in different shrimp organs are not just passive passengers; they form an interconnected and co-regulated team that actively influences the host’s phenotype and behavior. Even under changes in the host genotype, the same diet and environment can lead to changes in the shrimp tissue’s microbiota composition and function.

Moreover, specific microorganisms in the shrimp play crucial roles in lipid degradation, metabolic processes for short-chain fatty acids in the hepatopancreas, and glycan metabolism in the intestine (4). Therefore, by investigating the hologenome, integrating the host genetic makeup and their associated microbiota can offer valuable insights for developing new aquaculture strategies to enhance shrimp production, disease resistance, feed efficiency, and immune responses.

In this study, we addressed these critical questions by evaluating the structure and function of the shrimp microbiota in the hepatopancreas and intestine of two genetic lines raised under actual hatchery aquaculture conditions. One genetic line was chosen for its fast growth (Gen1), while the other was selected for its disease resistance (Gen2). Our goal was to understand the connection between host genetics and the microbiota in the two significant digestive organs of the shrimp. To do this, we used genotyping arrays with over 6,400 SNPs using the Illumina Infinium chip to assess the genetic variation of the shrimp. Additionally, we profiled the V3-V4 16S rRNA hypervariable regions. These analyses revealed the importance of host genetics in influencing the selection of taxa in the shrimp’s hepatopancreas and intestine in aquaculture conditions, offering potential for future applications in developing pre and probiotics to maintain the health and efficient production of this aquaculture species.

## Results

### 1. Experimental design under aquaculture conditions

The *Litopenaeus vannamei* were obtained from two production laboratories in Mexico, each with distinct phenotypic characteristics. One was a fast-growth (Gen 1), and the other was a disease-resistant (Gen 2) genetic line. All post-larvae were raised to an average weight of 12 ± 2 g in three independent aquaculture open ponds at the shrimp farm “Camarones el Renacimiento S.P.R. de R.I” in northern Mexico (Fig. 1). Two ponds (A and B) were in separated farm areas (Fig. 1). This setup allowed us to study how the microbiota structure was affected by different environments maintaining the same genetic line (Gen1). In contrast, Pond C, which was adjacent to Pond A, contained shrimps from a different genetic line (Gen2) to investigate the impact of two different genetic lines in a similar environment (Fig. 1). Importantly, all three ponds received the same aquaculture management techniques throughout the experiment (100 days).

**Figure 1.**
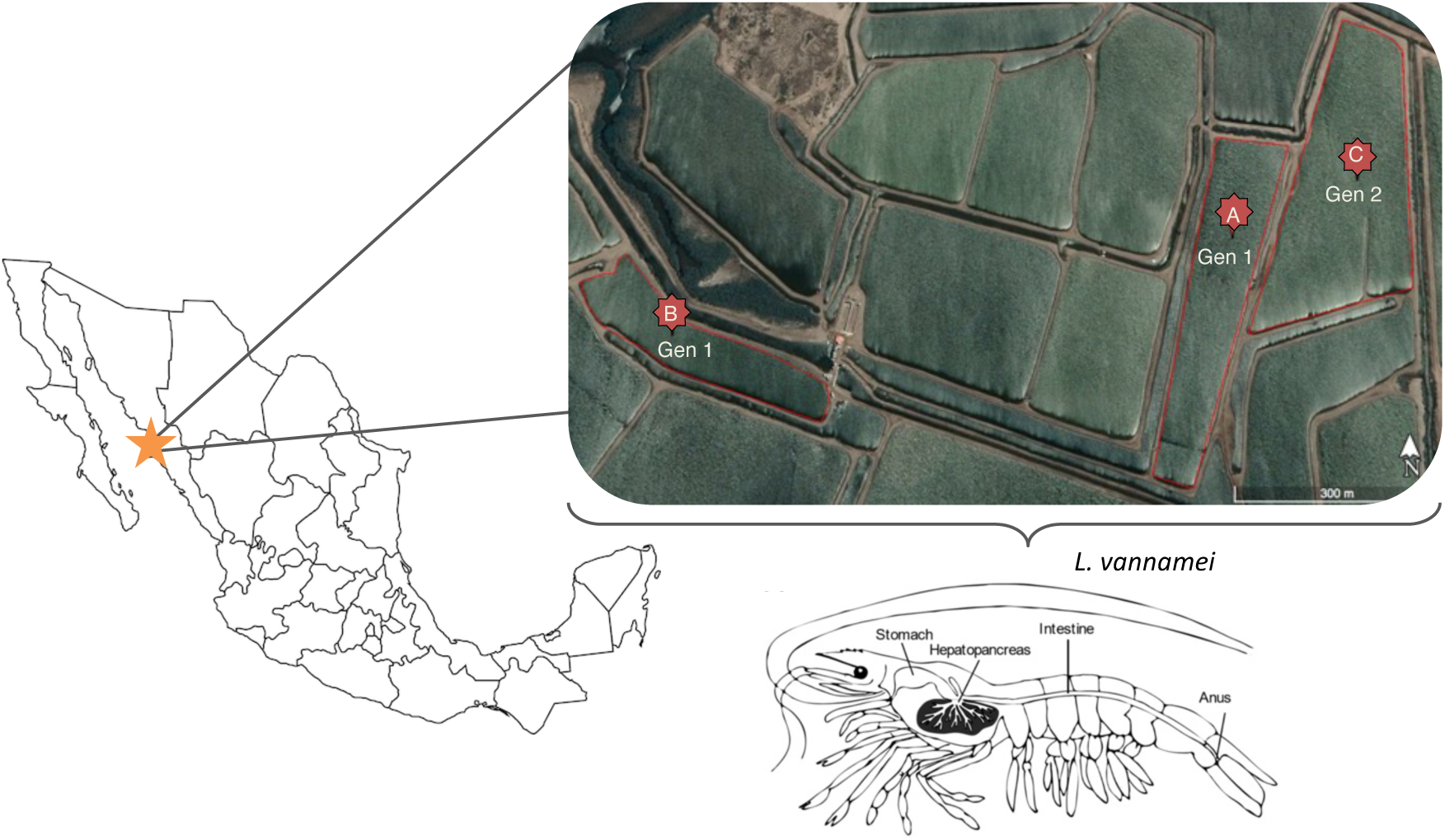
Experimental design for shrimp sample collection. Satellite overview of sample collection sites in Sinaloa, Mexico (CNES/Airbus©, 2020). Ponds A and B were located separated from each other and contained shrimps from the same genetic line (Gen1), while pond C was located next to pond A and contained shrimps from the second genetic line (Gen2). Below the map we show a diagram of the anatomical location of the dissected hepatopancreas and intestine.

### 2. Genome-wide SNP analysis confirmed two independent genetic lines

The post-larvae used in this study were sourced from two production laboratories, indicating they were from different genetic backgrounds due to distinct crossbreeding programs. To validate these genetic differences and establish both groups as independent genetic lines, we conducted genotyping using the Illumina Infinium ShrimpLD-24 array, which includes 6,465 SNPs, on 27 samples. Nine samples were taken from each pond.

After undergoing quality control, 4,476 SNPs with a call rate of 0.998 were selected for further study. We then conducted an ADMIXTURE analysis, which allows the classification of individuals into discrete subpopulations based on genetic diversity (21). This analysis was conducted for values of K ranging from 1 to 4, with the suitability of different K values assessed using cross-validation (CV). It was found that K=2 had the lowest CV error, assigning the 27 samples to two populations corresponding to the two genetic lines (Fig. 2A).

**Figure 2.**
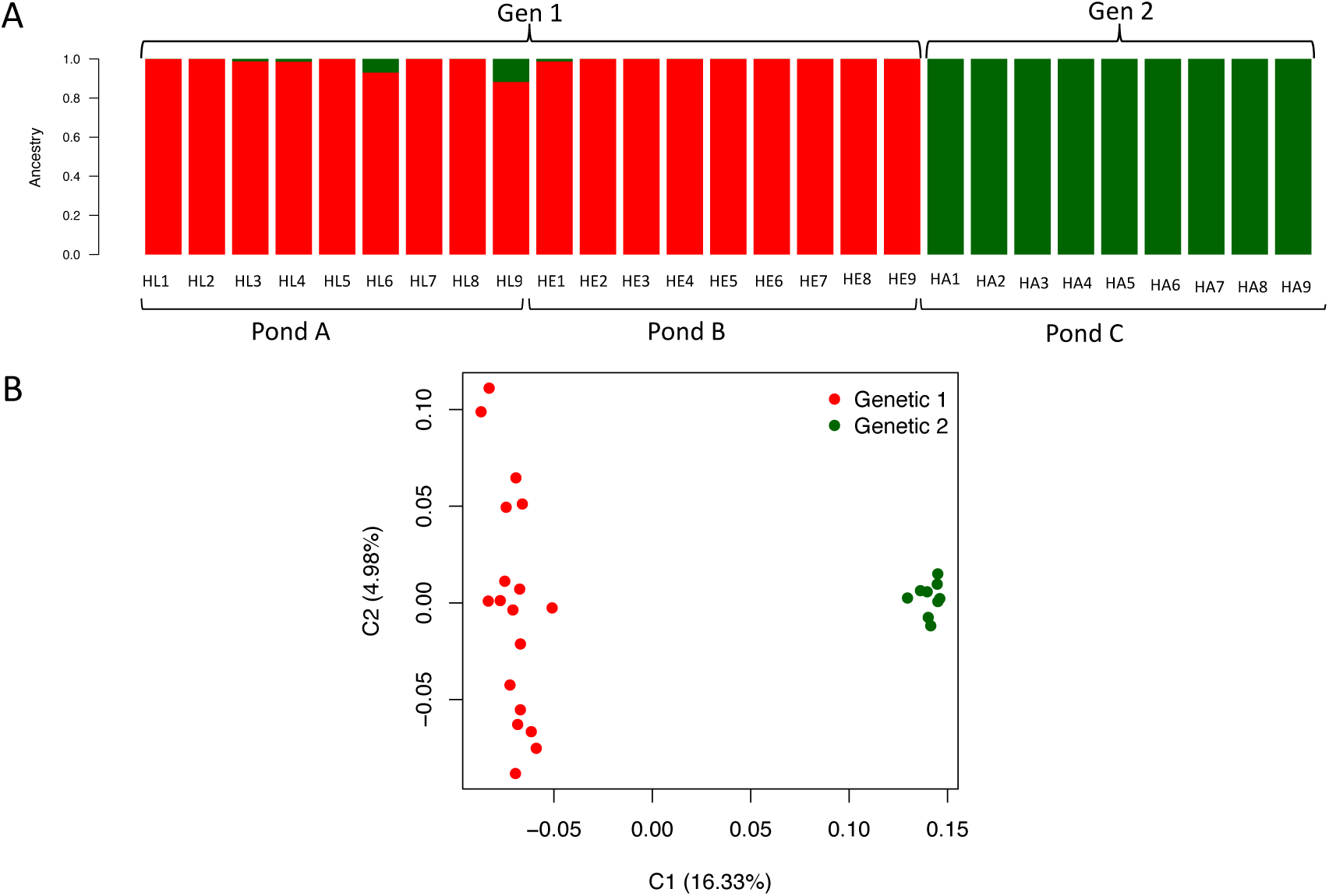
Genetic variability between both genetic lines. A) ADMIXTURE analysis (K= 2) of nine shrimp samples from each pond. The colors red and green indicate the two genetic populations represented in the samples. Samples from Ponds A and B correspond to the shrimps from Gen1, while samples from Pond C correspond to shrimps from Gen2. B) Multidimensional scaling analysis (MDS) with samples tagged by genetic line.

Furthermore, the multidimensional scaling analysis (MDS) revealed two clear and distinct clusters. One cluster included all samples from Gen 1 (ponds A and B), while the other cluster grouped all samples from Gen 2 (pond C) (Fig. 2B). These clusters were separated by the first principal component (C1) accounting for 16.33% of the total variability (Fig. 2B). Additionally, the estimated genetic differentiation (FST) between the two genetic lines was 0.174 (17.4%), indicating a significant difference between populations and confirming that Gen1 and Gen2 represented distinct and independent genetic lines. All these results confirmed that samples came from two different genetic populations (Figs. 2A and 2B).

### 3. The genetic line greatly influences the shrimp’s microbiota and growth performance

Microbial community profiles were obtained from sequencing of the V3-V4 hypervariable regions from the 16S ribosomal RNA (rRNA) gene (see Methods). Nine samples were taken from each pond. The ß-diversity (intersample diversity) results were observed in a principal coordinate analysis (PCoA) using UniFrac distances to analyze the differences. We then grouped the samples by organ (Fig. 3A) and genetic line (Fig. 3B). Additionally, we analyzed similarities (ANOSIM) and permutational analysis of variance (PERMANOVA) to explore the similarities between groups quantitatively.

**Figure 3.**
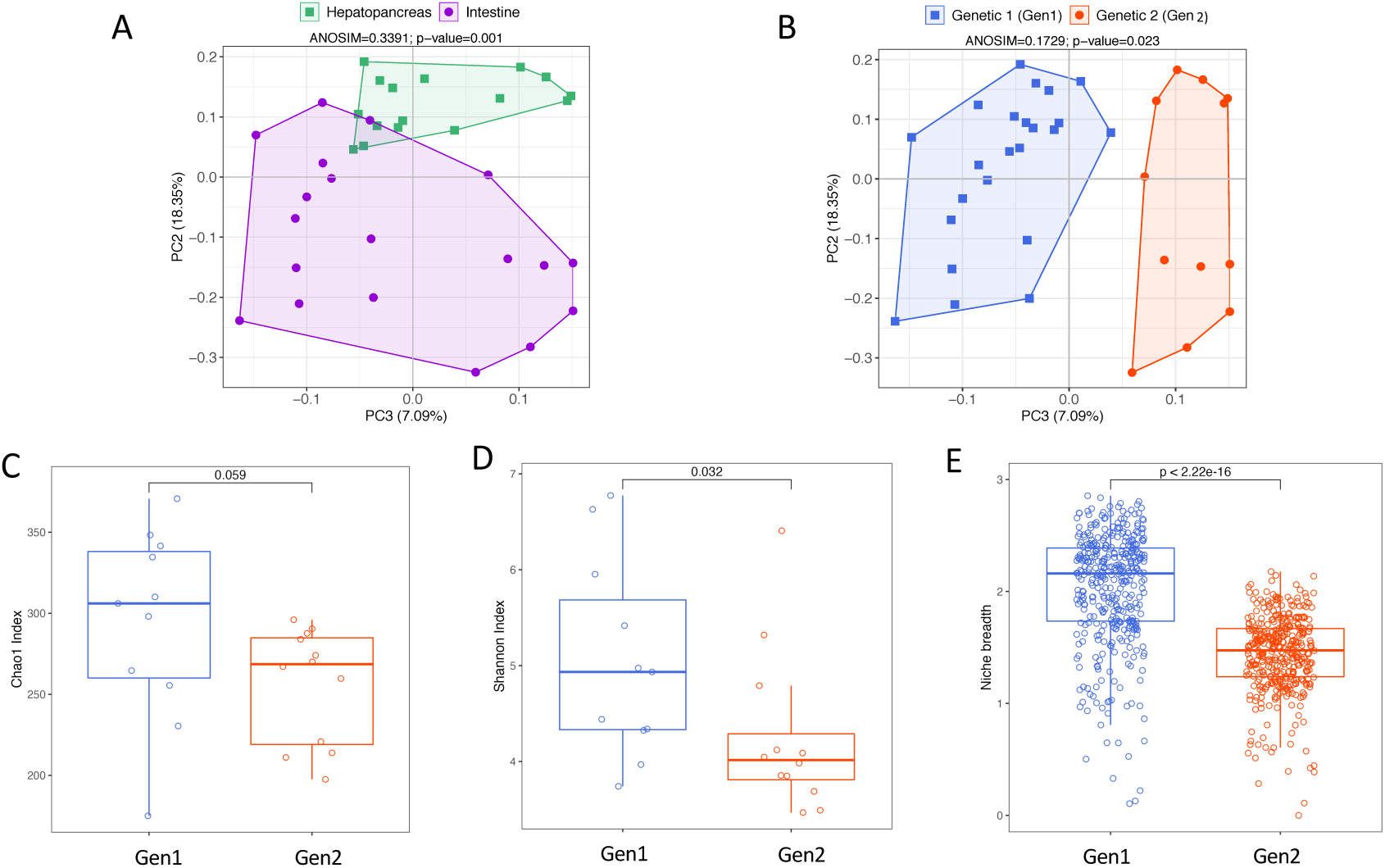
Alpha diversity indices and beta diversity analysis. Unweighted principal coordinate analysis (PCoA) of UniFrac distances representing the microbiota variability in samples tagged by A) organ, and B) genetic line. The ANOSIM R and p values are indicated above each graph. Boxplots showing the distribution for C) Chao1, and D) Shannon index between genetic lines. E) Niche breadth estimation for the microbiota in both genetics. All graphs consider both organs from pond A and C . Statistical differences between groups were evaluated with a Mann-Whitney test using a 95% confidence level of p<0.05.

As expected, upon categorizing the samples by organ (Fig. 3A), we observed significant (*p*<0.05) and the highest ANOSIM (R= 0.339, *p*= 0.001) and PERMANOVA (F= 6.05, *p*= 0.001) values, confirming that the organ was the most influential factor shaping the microbiota in the shrimp. Similarly, when we classified the samples by genetic line (Gen1 and Gen2), we observed a significant (p<0.05) albeit lower ANOSIM (R= 0.173, *p*= 0.023) and PERMANOVA (F= 2.483, *p*= 0.003) values, indicating that the genetic line was the second most influential factor modulating the shrimp microbiota (Fig. 3B).

In addition, to discard any potential impact of the pond on microbiota variability, we performed ß-diversity, ANOSIM, and PERMANOVA analyses only considering samples from ponds A and B, which were in distant areas within the farm but maintained the same genetic line (Fig. 1). The results showed low and nonsignificant values (p>0.05) (ANOSIM R=0.074, *p*=0.084; PERMANOVA F=1.834, *p*=0.069), suggesting that the pond did not influence the structure of the microbiota (Fig. S1). These findings further support our conclusion that the organ and the genetic line were the two major factors influencing the microbiota in our study.

### 4. The genetic line has a more relevant influence on shaping the microbiota in the hepatopancreas than in the intestine

To investigate how the host genetics affect the microbiota, we compared samples from ponds A and C, which were next to each other but housed shrimps from different genetic lines (Fig. 1). The overall diversity considering both organs, showed found that Gen1 had higher richness (Fig. 3C) and significantly (p<0.05) higher diversity (Fig. 3D) compared to Gen2. When we examined each organ separately, we consistently found higher richness and diversity in Gen1, although the difference was not statistically significant (Fig. S2). Notably, a niche-breadth estimation (Fig. 3E) revealed that the microbiota of Gen1 had a significantly wider niche breadth (p<0.0001) than Gen2, whether considering both organs together (Fig. 3E) or separately (Fig. S2). This suggests that the microbiota in Gen1 can exploit the available resources more efficiently than in Gen2.

The ANOSIM of Unifrac distances comparing the samples of Gen1 (pond A) versus Gen 2 (pond C) separating each organ revealed a significant (*p*= 0.026) effect of the genetic line on the microbial composition in the hepatopancreas, explaining approximately 30.0% (R=0.30) of the differences. This effect was also supported by PERMANOVA (F = 1.999; p = 0.044). However, in the intestine, there was no significant impact of the genetic line, as indicated by ANOSIM (R=0.122, p = 0.108) and PERMANOVA (F = 1.521 p = 0.097) analyses.

Furthermore, comparing operational taxonomic units (OTUs) between genetic lines for both organs revealed that Gen1 had more unique OTUs than Gen2, confirming the highest richness of Gen1 observed in alpha diversity analysis. Specifically, in the hepatopancreas, Gen1 had 9.5% unique OTUs, while Gen2 had 1%, and in the intestine, Gen1 had 1.5%, while Gen2 had 0% unique OTUs (Fig. S3). All these results suggest that the genetic line has a more significant impact on selecting the microbiota in the hepatopancreas than in the intestine.

Finally, we performed a LEfSe analysis to identify significantly enriched taxas in each genetic line. The results showed that there were 12 significantly enriched taxa for Gen1 and 19 for Gen2 in the hepatopancreas. In contrast, the intestine had 43 enriched taxa for Gen1 and 13 for Gen2 (Figs. 4A and B). Notably, Gen1 showed an enrichment of probiotic bacteria such as *Enterobacter, Ruegeria, Loktanella,* and *Brevibacteria*, while Gen2 presented an enrichment of some cyanobacteria such as *Arthrospira platensis, Nostocaceae, Lyngya,* and *Phormidium*.

**Figure 4.**
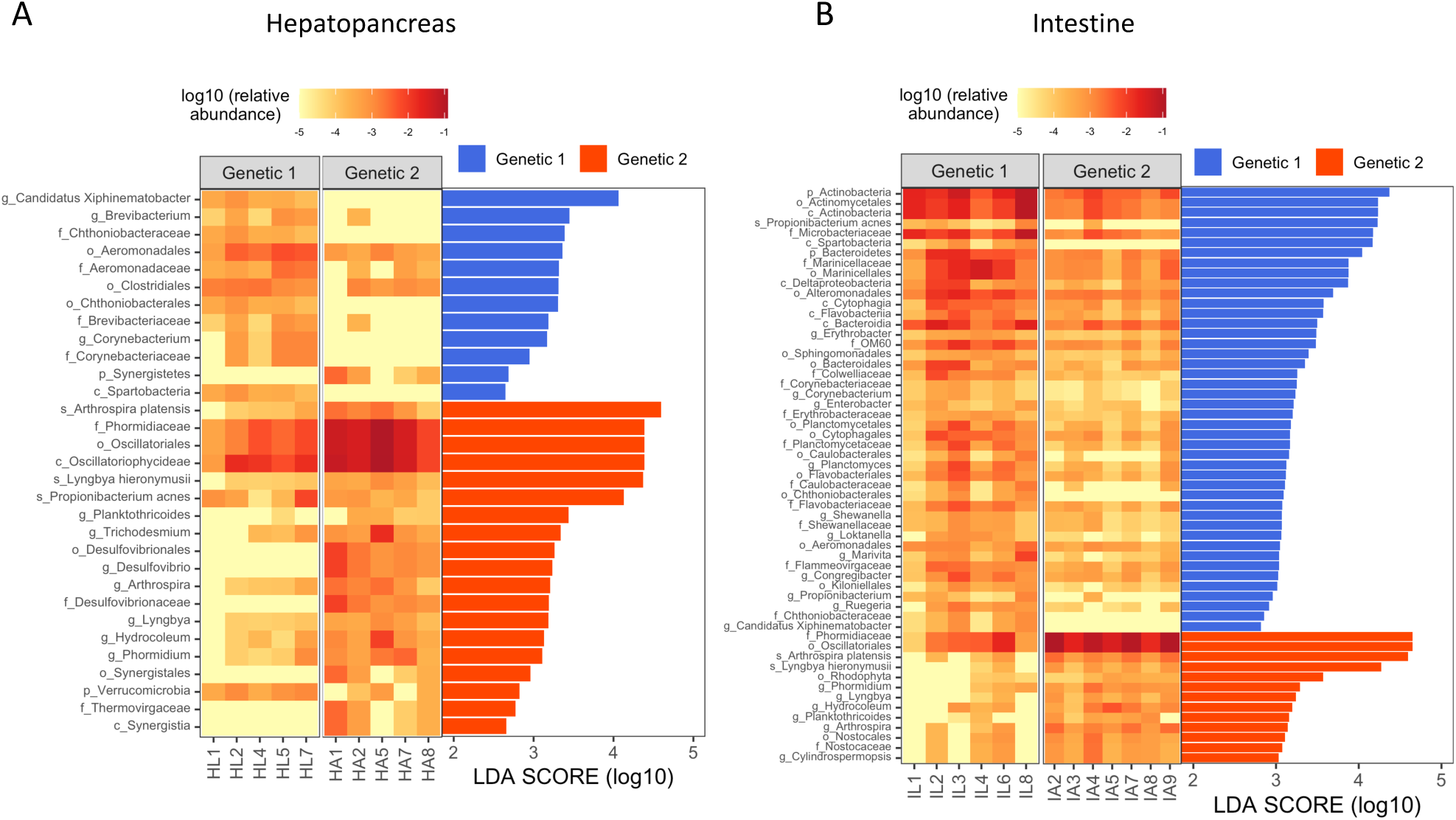
LEfSe analysis of enriched bacteria between both genetic lines. In A) hepatopancreas and B) intestine. The heat maps represents the relative abundance of each bacteria in all samples. The samples considered from Gen1 correspond to pond A, and samples from Gen2 correspond to pond C.

### 5. The genetic line influences the abundance of beneficial microbes in the shrimp microbiota

To explore if any of the genetic lines had an enrichment of beneficial microbes, we analyzed the abundance of 80 species proven to enhance shrimp health (22). We found that Gen1 had more of these beneficial microbes, with 19 species, while Gen2 had 17 species. Upon analyzing the abundance of each species, we discovered that Gen1 had a significant enrichment *(p* < 0.05) of *B. cereus* in the hepatopancreas

Interestingly, when we compared the abundance of each beneficial microbe between the two genetic lines, we found that only the intestine of Gen1 showed significant enrichment (p = 0.007) of beneficial microbes (Fig. 5B). In contrast, the differences in the hepatopancreas were not significant (p = 0.29) (Fig. 5A). These findings suggest that genetic lines may play a pivotal role in influencing the presence of beneficial microbes in the host’s microbiota, depending on the selected organ.

**Figure 5.**
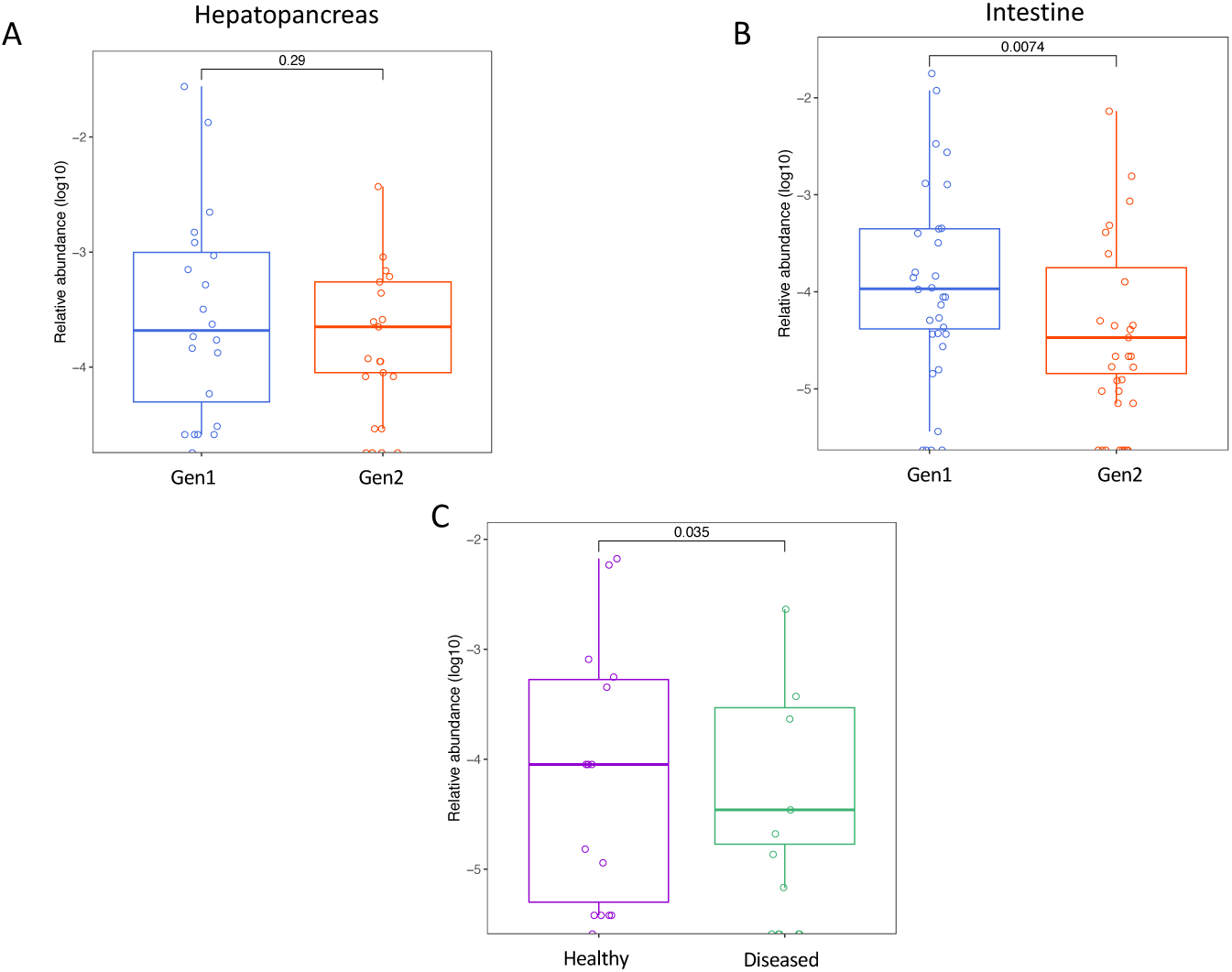
Distribution of the relative abundance of beneficial microbes. The boxplots compare the relative abundance (log10) of beneficial microbes between Gen1 and Gen2 at species level in A) hepatopancreas, B) intestine, and C) between the intestine of healthy and diseased shrimps. The statistical differences were determined with a Wilcoxon test (p<0.05).

### 6. The healthy status of shrimps influences the abundance of beneficial microbes

Our next step was to investigate the potential link between a higher presence of beneficial microbes and the overall health of shrimps. To this end, we analyzed the microbiota in the intestines of healthy and diseased (EMS-positive) shrimp samples that were previously collected by our research team (4). We then compared the abundance of beneficial microbes between the two groups. Our analysis showed that healthy shrimp showed a higher abundance of beneficial microbes than the diseased shrimp samples (Fig. S5). This suggests that a higher abundance of beneficial microbes in Gen1, as opposed to Gen2, could be a critical factor in maintaining a healthier status.

### 7. The wild-type genome and microbiota differs from the one established in both genetic lines

Previous observations in shrimp (4,10) and other organisms (23,24), including humans (25), suggest that native microbiota can be associated with better host fitness. To investigate whether the microbiota of the Gen1 and Gen2 resembled that of wild-type shrimp, we collected nine samples of wild-type shrimps, three of them were used for genotyping and nine for microbiota analyses. First, we confirmed that wild-type samples had a significantly different genetic structure than the two genetic lines (Figs. 6A and B). Second, we compared the wild-type microbiota with the genetic lines using an unweighted PCoA analysis with UniFrac distances and found two clusters along the X-axis, accounting for 24.64% of the variability (Fig. 6C). One cluster contained the samples from Gen1 and G2, while the second cluster grouped the wild-type samples. The ANOSIM analysis indicated a significant (*p* < 0.005) variability among these groups with an R-value of 0.548. These findings suggest that the microbial communities in the intestine microbiota of wild-type shrimp differ from those in the intestinal microbiota of both genetic lines and that neither of the two genetics is closer to the microbiota of wild shrimps.

**Figure 6.**
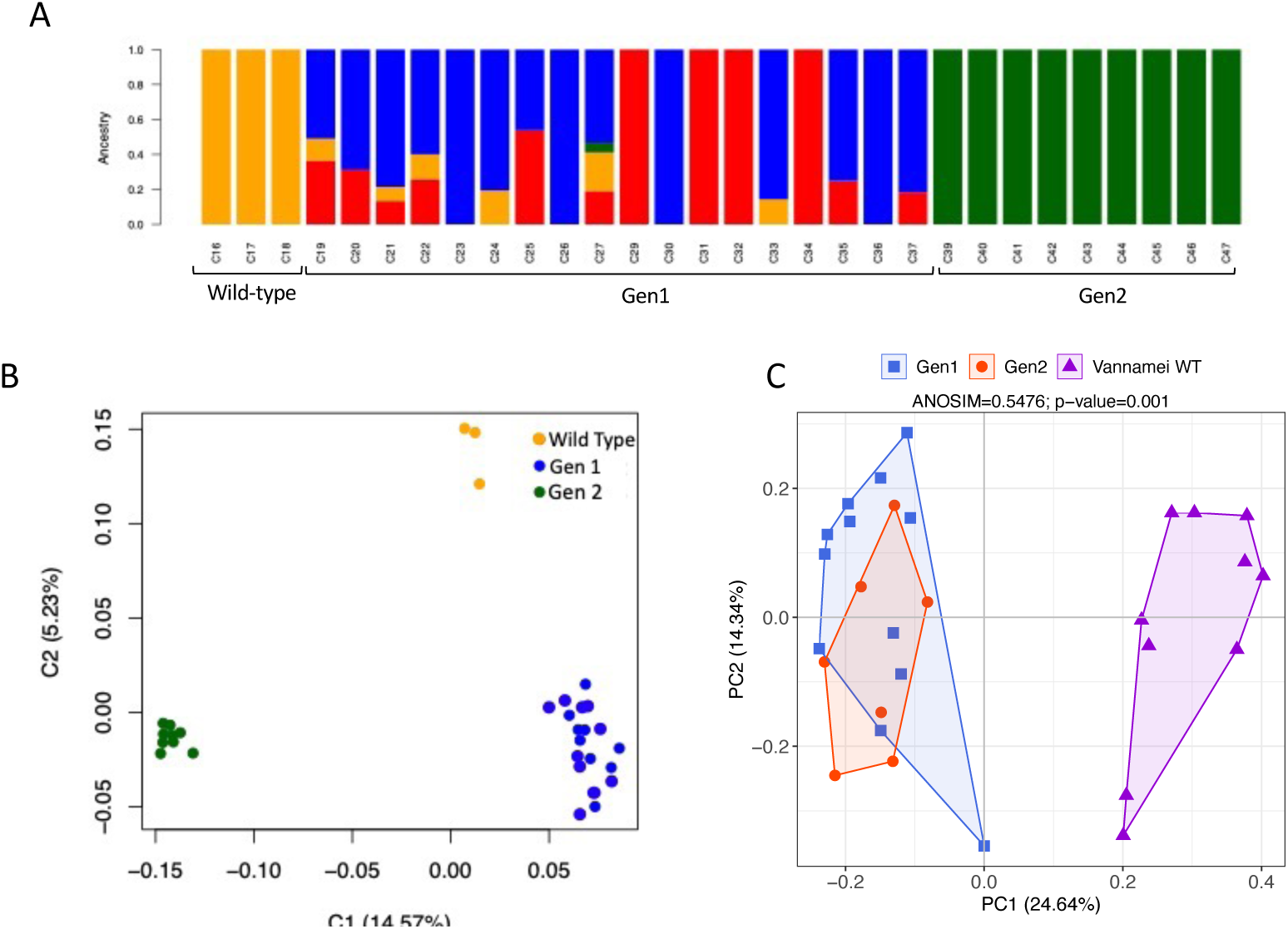
Genetic and microbiota variability between both genetic lines and Wild samples. A) ADMIXTURE analysis (K=4) of samples from wild-type, Gen1, and Gen2 origins. The colors yellow, blue, red and green indicate the genetic subpopulations represented in the samples. B) Multidimensional scaling analysis (MDS) representing the genetic variability in samples tagged by genetic line and wild-type origin. C) Unweighted principal coordinate analysis (PCoA) of UniFrac distances representig the microbiota variability with samples tagged by genetic line and wild-type origin. The ANOSIM R and p values are indicated above the graph.

## Discussion

The microbiota plays a pivotal role in maintaining health and promoting the development of any organism (26). It aids in the absorption of nutrients, regulation of metabolic processes, and modulation of immune responses (27,28). In shrimp, the dynamics of the microbiota are influenced by a variety of factors, both external and internal to the host, such as water quality (29), feeding formulations (9), diet probiotics (22), developmental stage (9), and diseases (4,7). Furthermore, genetic variations have been shown to impact the composition of these microbial communities, highlighting the crucial role of the host in shaping an effective microbiota (30,31).

The selection of genetic traits is crucial in all intensive agricultural activities, including animal husbandry. This recognition of the importance of genetic traits is also a cornerstone of the shrimp aquaculture industry. Many shrimp larvae production laboratories have developed crossbreeding programs to offer breeds with desirable characteristics for high-quality production, such as fast growth or disease resistance. However, combining all these traits in one “ideal larvae breed” has proven challenging. In this sense, understanding the microbiota and the influence of host genetics in shaping it could help us develop or complement these desirable traits.

To enhance our understanding, we conducted an analysis to determine whether there is a unique microbiota connected to two different genetic lines within the hepatopancreas and intestine of the same shrimp species (*L. vannamei*) under aquaculture hatchery conditions. We sought to investigate if this microbiota could indicate a superior fitness of one of the genetic lines. This finding could have far-reaching implications for aquaculture and genetic lines’ fitness.

The postlarvae used in the study were chosen based on their fast growth and disease resistance, two key traits in aquaculture. These postlarvae were sourced from two separate Mexican producers, each employing unique and intriguing crossbreeding strategies. Our research aimed to validate the significant genetic variability and distinct genetic lines of these selected strains. To achieve this, we conducted a genome microarray analysis, a comprehensive and detailed process. The admixture analysis indicated that the lowest cross-validation error occurred when K=2, and the FST value was 17.4% (0.174). In population studies in plants of the same species, F_ST_ values greater than 15% are deemed to indicate significant differentiation between populations, while values around 5% are considered insignificant (32). These results support the notion that genetic profiling in our study revealed two independent genetic backgrounds.

Extensive studies have shown that environmental conditions such as temperature, pH, salinity, and sediment organic compounds can impact the microbiota of shrimp (33,34). Thus, we maintained consistent hatchery conditions by applying the same technical management, feeding regime, and stocking density in the three studied ponds. Additionally, by raising the same genetic line in two different ponds (ponds A and B), beta analysis ruled out any influence of the pond on the microbiota composition.

Consistent with previous studies (9), our ANOSIM and Permanova analysis confirmed that the organ significantly shapes the microbiota composition, with the genetic line also playing a substantial role. Our data revealed that the genetic line has a more pronounced impact on the selection of the hepatopancreas microbiota, accounting for 30% of the variability, compared to the intestine’s 12%. These findings are supported by the LefSe analysis and the OTUs comparison in the Venn diagrams, which suggest that the hepatopancreas provides a more selective environment for microbial communities. This is likely due to the physiological function of the hepatopancreas as the main organ for enzyme activity, nutrient absorption, and immunity in crustaceans with less influence on the environment than the intestine (35), thereby imposing a stronger selective pressure on the colonizing bacteria. In contrast, the intestine appears to host a more adaptable microbiota that can respond to external influences, as seen in other studies (36).

The alpha diversity analysis revealed that Gen1 had higher microbial richness and significantly greater diversity than Gen2. This finding suggests potential implications for shrimp growth and ecosystem stability, which could greatly interest our field. In humans, a reduced diversity in the intestinal microbiota has been linked to diseases such as Crohn’s disease (37), irritable bowel syndrome (38) and colorectal cancer (39). On the other hand, a microbial community with high richness and diversity is considered more stable (40), resilient (41) and resistant to pathogen invasion (42,42,43). Additionally, the niche breadth estimation indicated a significantly wider niche breadth for Gen1 compared to Gen2. Previous studies suggest that a wider niche breadth corresponds to a generalist microbiota (44,45), while a microbiota with a narrow niche breadth is called a specialist (44,45). Therefore, the microbiota associated with Gen1 may exploit more diverse resources and tolerate a more variable environment than Gen2. Overall, the alpha diversity and niche breadth may point to a more complex bacterial ecosystem capable of more efficient nutrient and energy assimilation (46), which could contribute to better growth in the Fast-Growth shrimp strain.

Our research has revealed a significant difference in the enriched taxa between Gen1 and Gen2. In Gen1, we found taxa that are beneficial for their host’s metabolic health, such as the genus Brevibacteria, which produces short-chain fatty acids (47), and the genus Corynebacterium and class Bacteroida, which can enhance the immune system and metabolize dietary fibers (48,49). However, in Gen2, we observed an enrichment of diverse cyanobacteria, such as *Arthrospira platensis* (50), the family Nostocaceae and genera Lyngya and Phormidium (51–53). While these taxa have been associated with beneficial effects on human health, it is essential to note that some cyanobacteria, like *Nodular spumigena,* can pose a risk to shrimp aquaculture due to their potentially harmful impact on water quality and feed conversion ratios (54). Along these lines, the abundance analysis of microbes particularly beneficial for shrimp showed an increased abundance in Gen1, which could be associated with this genetic’s better growth performance. Additionally, there was a higher abundance of beneficial microbes in the microbiota of healthy shrimp than in EMS-positive shrimp. This suggests these bacteria are associated with better health status. The external administration of probiotics is a widely recommended strategy for diminishing diseases (5,55,55). However, their application in shrimp-rearing conditions has shown variable results and limited success (56–58). One potential explanation is that the applied probiotics cannot properly colonize and maintain viable populations (6,59). In this sense, identifying beneficial microbes that naturally reside in the shrimp microbiota could guide the design and administration of prebiotics to promote the proliferation of beneficial bacteria in the host. Furthermore, the finding that a specific genetic line of shrimp can increase the probiotics under aquaculture conditions opens the opportunity to be a selectable trait in the search for better shrimp larvae for production. Further, when considering that all the shrimp received the same technical management, it is worth noting that ponds with Gen1 had better productivity on the farm than Gen2. Specifically, at the end of the rearing cycle, Gen1 had a total production of 2.9 tons, while Gen2 had a total production of 2.3 tons. Interestingly, these results align with our findings that Gen1 had a microbiota with all the characteristics of being richer, more diverse, more resilient, and enriched in beneficial microbes.

There are several factors that impact the shrimp microbiota (9), with the surrounding sediment having the strongest influence (34) followed by water (60). Research on juvenile shrimps suggests that water can contribute as little as 8 % and as much as 29.69 % (60–62) to the shrimp microbiota. The sediment, on the other hand, is the major contributor, accounting for 30 % up to 60 % (63,64). Now, while further research is needed to quantify these contributions fully, our study suggests that genetic lineage contributes to approximately 17% of the microbiota variation.

Our research confirms the co-evolutionary relationship between the shrimp and its microbiota. It enhances our understanding of genetics as a factor influencing microbial communities and demonstrates its potential as a tool for intentional microbiome manipulation in aquaculture. It is evident that adopting a holobiome perspective is crucial for integrating characteristics such as genetics, development, environment, and microbiota into breeding and management programs, ultimately leading to improved growth efficiency, enhanced tolerance to environmental stress, and strengthened immune responses. Our next objective is to gather a more extensive dataset that will allow us to conduct genome-wide association studies (GWAS) aimed at identifying host genes associated with different microbiota profiles.

## Materials and methods

### Pond distribution and sample collection

The post-larvae of (*Litopenaeus vannamei)* from the fast-growth genetic line (Gen 1) were obtained from a de-identified reproduction laboratory in the South of Sinaloa, Mexico, and post-larvae from the disease-resistant genetic line (Gen 2) were obtained from a de-identified reproduction laboratory located in the North of Sinaloa, Mexico. All post-larvae were grown under the same conditions in the shrimp hatchery Camarones El Renacimiento located in the Northwest Mexican Pacific area of Sinaloa, Mexico (25°58′02.7″ N, 109°18′11.6″ W) (Fig. 1). The organisms were distributed in three ponds as follows: ponds A (26°01’49.3“N 109°23’21.9” W) and B (26°01’44.5“N 109°23’51.9” W) were located separately from each other and contained shrimps from Gen1 (Fig. 1), while pond C (26°01’55.4“N 109°23’12.5” W) was located next to pond A and contained shrimp from Gen2 (Fig. 1). All ponds received identical technical management (same water source, same feeding regime, same diet, same maintenance personnel, etc.). All ponds maintained similar conditions: an average water depth of 1.5 m, water salinity of approximately 40 ppm, average temperature of 29 °C, and a shrimp stocking density of 12 shrimps per square meter. All ponds were fed twice daily with standard commercial feed. After approximately three months of rearing, an average of six shrimp weighing 12 ± 2 g were collected from each pond. The hepatopancreas and intestine were aseptically dissected *in situ*, submerged individually in RNA-later® for 24 h at 4 °C (as recommended), and stored at -80°C until DNA extraction. In total, 35 samples (16 hepatopancreas and 19 intestines) were collected for this study.

### DNA extraction and amplicon sequencing

Total DNA was extracted from each organ using the Quick DNA Fecal/Soil Microbe Miniprep kit (Zymo Research, CA, USA, Cat. D6010) following the manufacturer’s recommendations. The concentration and integrity of the DNA were determined using a Qubit fluorometer (Life Technologies, CA, USA) and agarose gel electrophoresis. The 16S rRNA amplicons of the V3-V4 regions were generated as described in the 16S Metagenomics Sequencing Library Preparation user’s guide from Illumina (Illumina, CA, USA). All reactions were amplified under the following conditions: 95 °C for 3 min, 25 cycles of 95 °C for 30 s, 55 °C for 30 s, and 72 °C for 30 s, and a final elongation step at 72 °C for 5 min. After amplification, PCR amplicons were checked in 2% agarose gel, purified with Ampure XP beads (Beckman Coulter, Inc., CA, USA), and barcoded according to the 16S Metagenomics Sequencing Library Preparation user’s guide from Illumina. The quantity and size distribution of each library were assessed using the Qubit fluorometer and the Agilent 2100 Bioanalyzer (Agilent Technologies, CA, USA). All libraries were mixed in equal concentrations and sequenced on the Illumina Miseq sequencing platform using a 2×150 paired-end format at the National Institute of Genomic Medicine (INMEGEN) in Mexico City, México.

### Data preprocessing and taxonomic identification

The amplification primers for the corresponding 16S regions and Illumina adaptors were removed with Trimmomatic (v 0.4) (65). All trimmed sequences from each sample were joined (R1-R2) with fastq-join and filtered for quality (>Q20), where ambiguous bases were removed. The reads were clustered into 398 operational taxonomic units (OTUs) using QIIME (version 1.8) (66) and the uclust algorithm based on 3% divergence (97% sequence similarity) against the GreenGenes sequence database (version 13_5). The reverse strand matching option was enabled, and reads that did not hit against Green Genes were excluded. OUTs with an abundance of ≤ 0.01% were removed to exclude potential transitory microorganisms (67).

### Diversity (alpha and beta) and similarity (ANOSIM and Permanova) analyses

The alpha and beta diversity metrics were calculated using QIIME v1.9 from the final OTU table. The alpha diversity metrics were calculated at a sequence depth of 4,382 reads (the minimum number of reads obtained for the sequenced samples) and averaged from 10,000 iterations. The alpha diversity comparisons were evaluated using the Mann-Whitney test (nonparametric t-test) using a 95% level of confidence (*p*<0.05). The beta diversity was estimated by computing from the phylogenetic tree. The weighted and unweighted UniFrac distances among samples, and the UniFrac distance matrices were visualized using PCoA analysis. The robustness of the UPGMA tree was based on 1,000 replicates. To quantitatively assess the effects of different factors (pond, organ, and genetic line) on the microbiota, a permutational multivariate analysis of variance (PERMANOVA) (Anderson, 2001) with Adonis function was used on the unweighted UniFrac and Bray-Curtis distance matrices within QIIME. Additionally, the difference among groups in the distance matrices was evaluated with ANOSIM for every beta analysis.

### Linear Discriminant (LEfSe), niche breadth analyses and beneficial microbes analyses

For the differential abundance analysis, OTUs between ponds and host genetics were compared using LefSe. A significance level (alpha) of 0.05 and an LDA threshold >2 were applied to identify significantly different taxas. OTUs were subjected to a differential abundance analysis with LefSe, considering a significance level (alpha) of 0.05 and an LDA threshold >2. The niche breadth estimation was calculated using the niche.width function of the spa R package; the final results were plotted with ggplot2 in R. Further, the beneficial microbes were identified following the previously reported method (22) and the statistical differences between groups were calculated with a Wilcoxon test (p value ≤ 0.05).

### Genotyping arrays

A total of 30 genomic DNA samples (9 from each pond plus three blind samples duplicated for quality control) were genotyped using the Illumina Infinium ShrimpLD-24 v1.0 beadchip genotyping array, which included 6,465 SNPs (68). Quality control of the genotype data involved removing SNPs with a minor allele frequency of less than 0.05 and a call rate of less than 0.95. Pairwise identity-by-descent (IBD) estimates were used to identify the cryptic relationships among samples. The genotype data from the 27 samples were used for population admixture analysis. A total of 4,476 SNPs were used to perform a multidimensional scaling analysis (MDS) based on pairwise IBD estimates using PLINK (69). ADMIXTURE estimated individual ancestry proportions for K = 1 to K = 4. The fit of different K values was assessed using cross-validation (CV) procedures, where K = 2 showed the lowest CV error.

## Acknowledgments

We thank M.T.I Juan Manuel Hurtado Ramírez for informatics technical support as well as Bio. Filiberto Sánchez López for experimental and technical support. M.C. Alfredo Mendoza-Vargas from Unidad de Secuenciación Masiva and M.C. Raúl Mojica Espinosa from the Unidad de Microarreglos from the National Institute of Genomic Medicine (INMEGEN). Also the authors would like to thank the “Unidad Universitaria de Secuenciación Masiva y Bioinformática” of the “Laboratorio Nacional de Apoyo Tecnológico a las Ciencias Genómicas,” UNAM, especially Lizeth A. Matías Valdez for the technical sequencing support. And to the shrimp farm Camarones el Renacimiento S.P.R. de R.I. for providing the infrastructure and technical support during the sample collection.

## Author contributions

Experimental design and conceptualization F.C.G, L.G.B and A.O.L, sample collection A.L.Z. and A.C.H, data curation L.G.B and A.O.L, formal analysis L.G.B, S.R.H, F.C.G. and A.O.L, investigation and methodology F.C.G, L.G.B, R.S.M and A.O.L funding acquisition F.C.G and A.O.L, writing original draft F.C.G, R.S.M and A.O.L, writing, review and editing F.C.G, and A.O.L

## Data availability

The sequencing data relevant to this publication is available at NCBI under BioProject PRJNA1153518.

## Ethics

An ethics statement was not required for the current study as locations for the specimen collection are not protected, and field studies did not include endangered or protected species. Animals were sacrificed under university protocols to avoid animal suffering.

## References

1. Nicholson JK, Holmes E, Kinross J, Burcelin R, Gibson G, Jia W, et al. Host-Gut Microbiota Metabolic Interactions. Science. 2012 Jun 8;336(6086):1262–7.

2. The State of World Fisheries and Aquaculture 2020 [Internet]. FAO; 2020 [cited 2024 Sep 9]. Available from: http://www.fao.org/documents/card/en/c/ca9229en

3. Zhang W, Belton B, Edwards P, Henriksson PJG, Little DC, Newton R, et al. Aquaculture will continue to depend more on land than sea. Nature. 2022 Mar 10;603(7900):E2–4.

4. Cornejo-Granados F, Lopez-Zavala AA, Gallardo-Becerra L, Mendoza-Vargas A, Sánchez F, Vichido R, et al. Microbiome of Pacific Whiteleg shrimp reveals differential bacterial community composition between Wild, Aquacultured and AHPND/EMS outbreak conditions. Sci Rep. 2017 Dec;7(1):11783.

5. Li E, Xu C, Wang X, Wang S, Zhao Q, Zhang M, et al. Gut Microbiota and its Modulation for Healthy Farming of Pacific White Shrimp *Litopenaeus vannamei*. Rev Fish Sci Aquac. 2018 Jul 3;26(3):381–99.

6. Xiong J. Progress in the gut microbiota in exploring shrimp disease pathogenesis and incidence. Appl Microbiol Biotechnol. 2018 Sep;102(17):7343–50.

7. Xiong J, Zhu J, Dai W, Dong C, Qiu Q, Li C. Integrating gut microbiota immaturity and disease-discriminatory taxa to diagnose the initiation and severity of shrimp disease. Environ Microbiol. 2017 Apr;19(4):1490–501.

8. MacLeod MJ, Hasan MR, Robb DHF, Mamun-Ur-Rashid M. Quantifying greenhouse gas emissions from global aquaculture. Sci Rep. 2020 Jul 15;10(1):11679.

9. Cornejo-Granados F, Gallardo-Becerra L, Leonardo-Reza M, Ochoa-Romo JP, Ochoa-Leyva A. A meta-analysis reveals the environmental and host factors shaping the structure and function of the shrimp microbiota. PeerJ. 2018 Aug 10;6:e5382.

10. Rungrassamee W, Klanchui A, Chaiyapechara S, Maibunkaew S, Tangphatsornruang S, Jiravanichpaisal P, et al. Bacterial Population in Intestines of the Black Tiger Shrimp (Penaeus monodon) under Different Growth Stages. Bereswill S, editor. PLoS ONE. 2013 Apr 5;8(4):e60802.

11. Rahman M, Sabir AA, Mukta JA, Khan MdMA, Mohi-Ud-Din M, Miah MdG, et al. Plant probiotic bacteria Bacillus and Paraburkholderia improve growth, yield and content of antioxidants in strawberry fruit. Sci Rep. 2018 Feb 6;8(1):2504.

12. Luise D, Bosi P, Raff L, Amatucci L, Virdis S, Trevisi P. Bacillus spp. Probiotic Strains as a Potential Tool for Limiting the Use of Antibiotics, and Improving the Growth and Health of Pigs and Chickens. Front Microbiol. 2022 Feb 7;13:801827.

13. El-Saadony MT, Alagawany M, Patra AK, Kar I, Tiwari R, Dawood MAO, et al. The functionality of probiotics in aquaculture: An overview. Fish Shellfish Immunol. 2021 Oct;117:36–52.

14. Kostic AD, Howitt MR, Garrett WS. Exploring host–microbiota interactions in animal models and humans. Genes Dev. 2013 Apr 1;27(7):701–18.

15. Landsman A, St-Pierre B, Rosales-Leija M, Brown M, Gibbons W. Investigation of the Potential Effects of Host Genetics and Probiotic Treatment on the Gut Bacterial Community Composition of Aquaculture-raised Pacific Whiteleg Shrimp, Litopenaeus vannamei. Microorganisms. 2019 Jul 26;7(8):217.

16. Malard F, Dore J, Gaugler B, Mohty M. Introduction to host microbiome symbiosis in health and disease. Mucosal Immunol. 2021 May;14(3):547–54.

17. Soldan R, Fusi M, Cardinale M, Daffonchio D, Preston GM. The effect of plant domestication on host control of the microbiota. Commun Biol. 2021 Aug 5;4(1):936.

18. Maraci Ö, Antonatou-Papaioannou A, Jünemann S, Castillo-Gutiérrez O, Busche T, Kalinowski J, et al. The Gut Microbial Composition Is Species-Specific and Individual-Specific in Two Species of Estrildid Finches, the Bengalese Finch and the Zebra Finch. Front Microbiol. 2021 Feb 19;12:619141.

19. Tkacz A, Bestion E, Bo Z, Hortala M, Poole PS. Influence of Plant Fraction, Soil, and Plant Species on Microbiota: a Multikingdom Comparison. Bailey MJ, editor. mBio. 2020 Feb 25;11(1):e02785–19.

20. Zilber-Rosenberg I, Rosenberg E. Role of microorganisms in the evolution of animals and plants: the hologenome theory of evolution. FEMS Microbiol Rev. 2008 Aug;32(5):723–35.

21. Holsinger KE, Weir BS. Genetics in geographically structured populations: defining, estimating and interpreting FST. Nat Rev Genet. 2009 Sep;10(9):639–50.

22. Ochoa-Romo JP, Cornejo-Granados F, Lopez-Zavala AA, Viana MT, Sánchez F, Gallardo-Becerra L, et al. Agavin induces beneficial microbes in the shrimp microbiota under farming conditions. Sci Rep. 2022 Apr 16;12(1):6392.

23. Gazzaniga FS, Kasper DL. Wild gut microbiota protects from disease. Cell Res. 2018 Feb;28(2):135–6.

24. Rosshart SP, Vassallo BG, Angeletti D, Hutchinson DS, Morgan AP, Takeda K, et al. Wild Mouse Gut Microbiota Promotes Host Fitness and Improves Disease Resistance. Cell. 2017 Nov;171(5):1015–1028.e13.

25. Schauer DB. Indigenous microflora: Paving the way for pathogens? Curr Biol. 1997 Feb;7(2):R75–7.

26. Lee JY, Tsolis RM, Bäumler AJ. The microbiome and gut homeostasis. Science. 2022 Jul;377(6601):eabp9960.

27. Pickard JM, Zeng MY, Caruso R, Núñez G. Gut microbiota: Role in pathogen colonization, immune responses, and inflammatory disease. Immunol Rev. 2017 Sep;279(1):70–89.

28. Rooks MG, Garrett WS. Gut microbiota, metabolites and host immunity. Nat Rev Immunol. 2016 Jun;16(6):341–52.

29. Huang F, Pan L, Song M, Tian C, Gao S. Microbiota assemblages of water, sediment, and intestine and their associations with environmental factors and shrimp physiological health. Appl Microbiol Biotechnol. 2018 Oct;102(19):8585–98.

30. Bonder MJ, Kurilshikov A, Tigchelaar EF, Mujagic Z, Imhann F, Vila AV, et al. The effect of host genetics on the gut microbiome. Nat Genet. 2016 Nov;48(11):1407–12.

31. Goodrich JK, Waters JL, Poole AC, Sutter JL, Koren O, Blekhman R, et al. Human Genetics Shape the Gut Microbiome. Cell. 2014 Nov;159(4):789–99.

32. Frankham R, Ballou JD, Briscoe DA, McInnes KH. Introduction to Conservation Genetics [Internet]. 1st ed. Cambridge University Press; 2002 [cited 2024 Sep 9]. Available from: https://www.cambridge.org/core/product/identifier/9780511808999/type/book

33. Fan L, Wang Z, Chen M, Qu Y, Li J, Zhou A, et al. Microbiota comparison of Pacific white shrimp intestine and sediment at freshwater and marine cultured environment. Sci Total Environ. 2019 Mar;657:1194–204.

34. Zhang M, Pan L, Huang F, Gao S, Su C, Zhang M, et al. Metagenomic analysis of composition, function and cycling processes of microbial community in water, sediment and effluent of Litopenaeus vannamei farming environments under different culture modes. Aquaculture. 2019 May;506:280–93.

35. Vogt G. Functional cytology of the hepatopancreas of decapod crustaceans. J Morphol. 2019 Jul 12;jmor.21040.

36. Chen CY, Chen PC, Weng FCH, Shaw GTW, Wang D. Habitat and indigenous gut microbes contribute to the plasticity of gut microbiome in oriental river prawn during rapid environmental change. Jadhao SB, editor. PLOS ONE. 2017 Jul 17;12(7):e0181427.

37. Matsuoka K, Kanai T. The gut microbiota and inflammatory bowel disease. Semin Immunopathol. 2015 Jan;37(1):47–55.

38. Sha S, Xu B, Wang X, Zhang Y, Wang H, Kong X, et al. The biodiversity and composition of the dominant fecal microbiota in patients with inflammatory bowel disease. Diagn Microbiol Infect Dis. 2013 Mar;75(3):245–51.

39. Ahn J, Sinha R, Pei Z, Dominianni C, Wu J, Shi J, et al. Human Gut Microbiome and Risk for Colorectal Cancer. JNCI J Natl Cancer Inst. 2013 Dec 18;105(24):1907–11.

40. Larsen OFA, Claassen E. The mechanistic link between health and gut microbiota diversity. Sci Rep. 2018 Feb 1;8(1):2183.

41. Lozupone CA, Stombaugh JI, Gordon JI, Jansson JK, Knight R. Diversity, stability and resilience of the human gut microbiota. Nature. 2012 Sep;489(7415):220–30.

42. Lin L, Zhang J. Role of intestinal microbiota and metabolites on gut homeostasis and human diseases. BMC Immunol. 2017 Dec;18(1):2.

43. Tap J, Furet J, Bensaada M, Philippe C, Roth H, Rabot S, et al. Gut microbiota richness promotes its stability upon increased dietary fibre intake in healthy adults. Environ Microbiol. 2015 Dec;17(12):4954–64.

44. Deines P, Hammerschmidt K, Bosch TCG. Exploring the Niche Concept in a Simple Metaorganism. Front Microbiol. 2020 Aug 11;11:1942.

45. Luan L, Liang C, Chen L, Wang H, Xu Q, Jiang Y, et al. Coupling Bacterial Community Assembly to Microbial Metabolism across Soil Profiles. Bouskill N, editor. mSystems. 2020 Jun 30;5(3):10.1128/msystems.00298-20.

46. Hosomi K, Kunisawa J. Diversity of energy metabolism in immune responses regulated by micro-organisms and dietary nutrition. Int Immunol. 2020 Jun 26;32(7):447–54.

47. Murakami A, Toyomoto K, Namai F, Sato T, Fujii T, Tochio T, et al. Oral administration of *BREVIBACTERIUM LINENS* from washed cheese increases the proportions of short-chain fatty acid-producing bacteria and lactobacilli in the gut microbiota of mice. Anim Sci J. 2023 Jan;94(1):e13905.

48. Menberu MA, Liu S, Cooksley C, Hayes AJ, Psaltis AJ, Wormald PJ, et al. Corynebacterium accolens Has Antimicrobial Activity against Staphylococcus aureus and Methicillin-Resistant S. aureus Pathogens Isolated from the Sinonasal Niche of Chronic Rhinosinusitis Patients. Pathogens. 2021 Feb 14;10(2):207.

49. Zafar H, Saier MH. Gut *Bacteroides* species in health and disease. Gut Microbes. 2021 Jan 1;13(1):1848158.

50. Gentscheva G, Nikolova K, Panayotova V, Peycheva K, Makedonski L, Slavov P, et al. Application of Arthrospira platensis for Medicinal Purposes and the Food Industry: A Review of the Literature. Life. 2023 Mar 21;13(3):845.

51. Swain S, Bej S, Bishoyi AK, Mandhata CP, Sahoo CR, Padhy RN. Recent progression on phytochemicals and pharmacological properties of the filamentous cyanobacterium Lyngbya sp. Naunyn Schmiedebergs Arch Pharmacol. 2023 Oct;396(10):2197–216.

52. Tena Pérez V, Apaza Ticona L, Cabanillas AH, Maderuelo Corral S, Rosero Valencia DF, Quintana AM, et al. Anti-inflammatory activity of glycolipids isolated from cyanobacterium *Nodularia harveyana*. Nat Prod Res. 2021 Dec 17;35(24):6204–9.

53. Zampieri RM, Adessi A, Caldara F, Codato A, Furlan M, Rampazzo C, et al. Anti-Inflammatory Activity of Exopolysaccharides from Phormidium sp. ETS05, the Most Abundant Cyanobacterium of the Therapeutic Euganean Thermal Muds, Using the Zebrafish Model. Biomolecules. 2020 Apr 10;10(4):582.

54. Duan Y, Xing Y, Huang J, Nan Y, Li H, Dong H. Toxicological response of Pacific white shrimp Litopenaeus vannamei to a hazardous cyanotoxin nodularin exposure. Environ Pollut. 2023 Feb;318:120950.

55. Ninawe AS, Selvin J. Probiotics in shrimp aquaculture: Avenues and challenges. Crit Rev Microbiol. 2009 Feb;35(1):43–66.

56. Adel M, Yeganeh S, Dawood MAO, Safari R, Radhakrishnan S. Effects of *Pediococcus pentosaceus* supplementation on growth performance, intestinal microflora and disease resistance of white shrimp, *Litopenaeus vannamei*. Aquac Nutr. 2017 Dec;23(6):1401–9.

57. Liu KF, Chiu CH, Shiu YL, Cheng W, Liu CH. Effects of the probiotic, Bacillus subtilis E20, on the survival, development, stress tolerance, and immune status of white shrimp, Litopenaeus vannamei larvae. Fish Shellfish Immunol. 2010 May;28(5–6):837–44.

58. Xiong J, Dai W, Li C. Advances, challenges, and directions in shrimp disease control: the guidelines from an ecological perspective. Appl Microbiol Biotechnol. 2016 Aug;100(16):6947–54.

59. Giatsis C, Sipkema D, Ramiro-Garcia J, Bacanu GM, Abernathy J, Verreth J, et al. Probiotic legacy effects on gut microbial assembly in tilapia larvae. Sci Rep. 2016 Sep 27;6(1):33965.

60. Xiong J, Xuan L, Yu W, Zhu J, Qiu Q, Chen J. Spatiotemporal successions of shrimp gut microbial colonization: high consistency despite distinct species pool. Environ Microbiol. 2019 Apr;21(4):1383–94.

61. Li H, Gu S, Wang L, Shi W, Jiang Q, Wan X. Dynamic Changes of Environment and Gut Microbial Community of Litopenaeus vannamei in Greenhouse Farming and Potential Mechanism of Gut Microbial Community Construction. Fishes. 2024 Apr 26;9(5):155.

62. Xiong J, Dai W, Qiu Q, Zhu J, Yang W, Li C. Response of host–bacterial colonization in shrimp to developmental stage, environment and disease. Mol Ecol. 2018 Sep;27(18):3686–99.

63. Huang Z, Hou D, Zhou R, Zeng S, Xing C, Wei D, et al. Environmental Water and Sediment Microbial Communities Shape Intestine Microbiota for Host Health: The Central Dogma in an Anthropogenic Aquaculture Ecosystem. Front Microbiol. 2021 Nov 2;12:772149.

64. Zhang X, Li X, Lu J, Qiu Q, Chen J, Xiong J. Quantifying the importance of external and internal sources to the gut microbiota in juvenile and adult shrimp. Aquaculture. 2021 Jan;531:735910.

65. Bolger AM, Lohse M, Usadel B. Trimmomatic: a flexible trimmer for Illumina sequence data. Bioinformatics. 2014 Aug 1;30(15):2114–20.

66. Caporaso JG, Kuczynski J, Stombaugh J, Bittinger K, Bushman FD, Costello EK, et al. QIIME allows analysis of high-throughput community sequencing data. Nat Methods. 2010 May;7(5):335–6.

67. Bokulich NA, Subramanian S, Faith JJ, Gevers D, Gordon JI, Knight R, et al. Quality-filtering vastly improves diversity estimates from Illumina amplicon sequencing. Nat Methods. 2013 Jan;10(1):57–9.

68. Jones DB, Jerry DR, Khatkar MS, Raadsma HW, Steen HVD, Prochaska J, et al. A comparative integrated gene-based linkage and locus ordering by linkage disequilibrium map for the Pacific white shrimp, Litopenaeus vannamei. Sci Rep. 2017 Sep 4;7(1):10360.

69. Purcell S, Neale B, Todd-Brown K, Thomas L, Ferreira MAR, Bender D, et al. PLINK: A Tool Set for Whole-Genome Association and Population-Based Linkage Analyses. Am J Hum Genet. 2007 Sep;81(3):559–75.

