## Supplementary Figures for "Host Genome Drives the Diversity, Richness, and Beneficial Microbes in the Shrimp Microbiota: A Hologenome Perspective"

### SUPPLEMENTARY MATERIAL

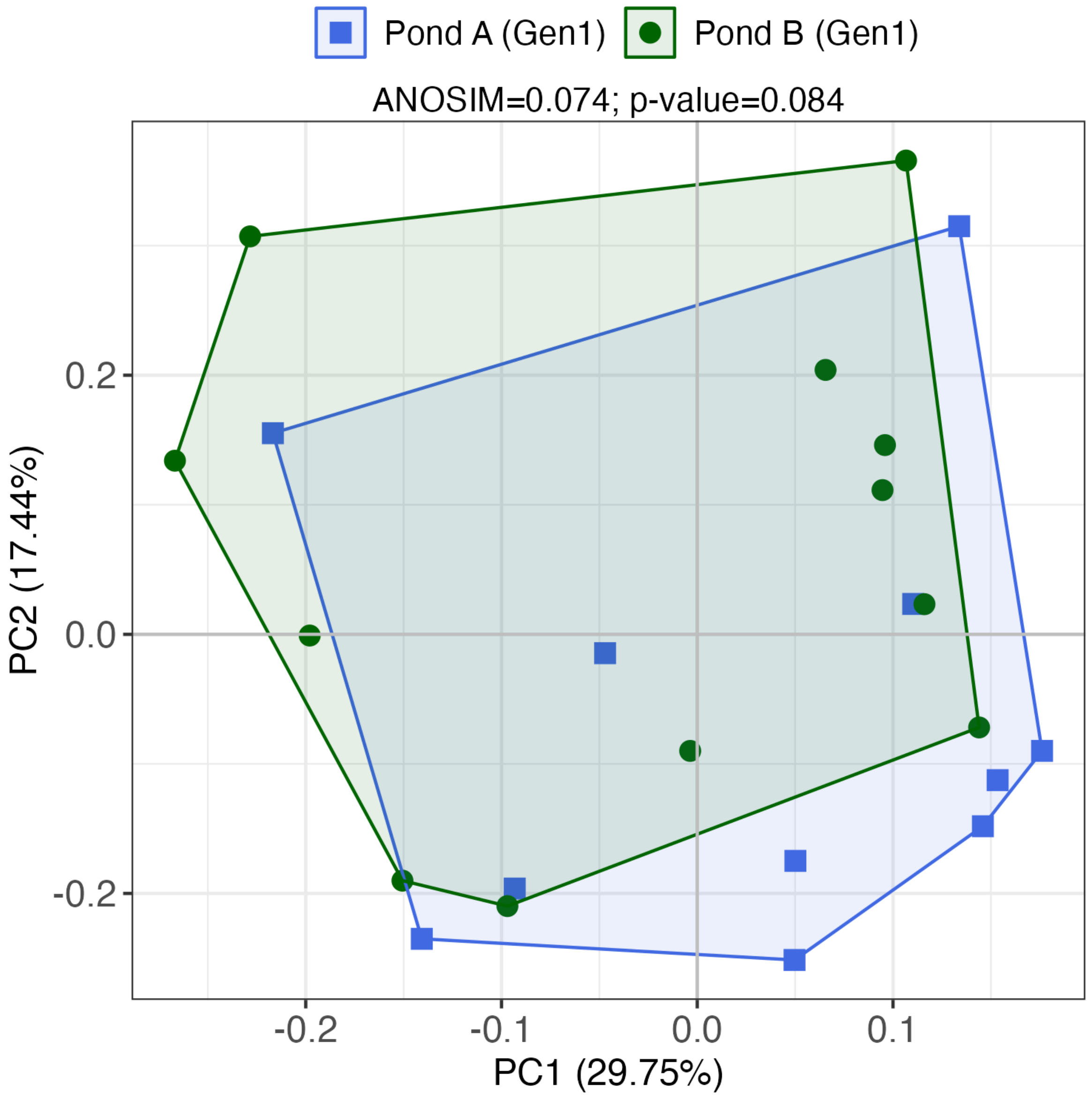


**Supplementary Figure 1. Beta diversity analysis.** Unweighted principal coordinate analysis (PCoA) of UniFrac distances representing the microbiota variability in samples tagged by pond. The ANOSIM R and p values are indicated above the graph.

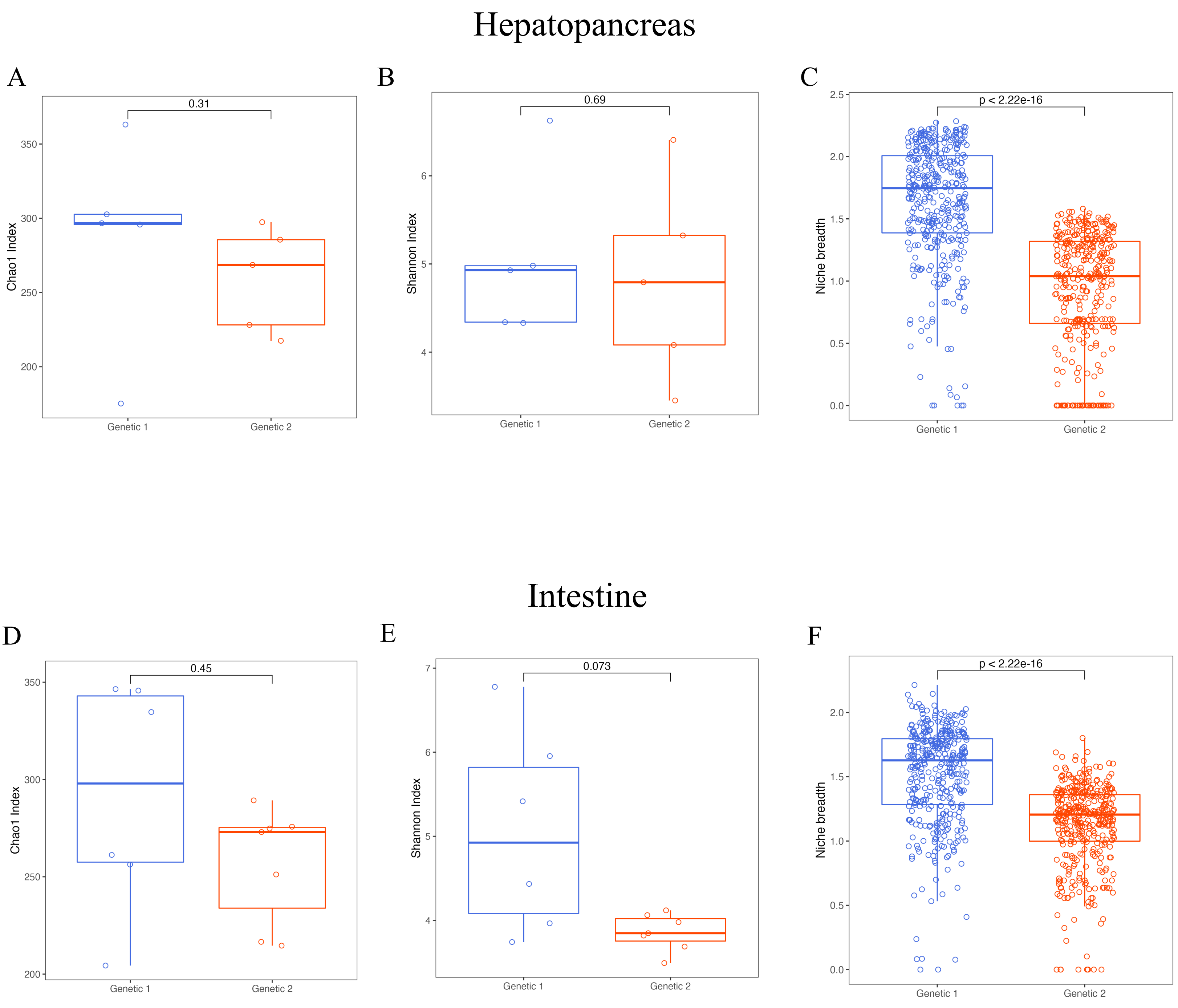


**Supplementary Figure 2. Alpha diversity indices and niche breadth estimation.** Boxplots showing the richness, diversity and niche breadth distribution of the microbiota in both genetic lines considering the hepatopancreas (panels A, B and C), and the intestine (panels D, E and F). Statistical differences between groups were evaluated with a Mann-Whitney test using a 95% confidence level of $p<0.05$.

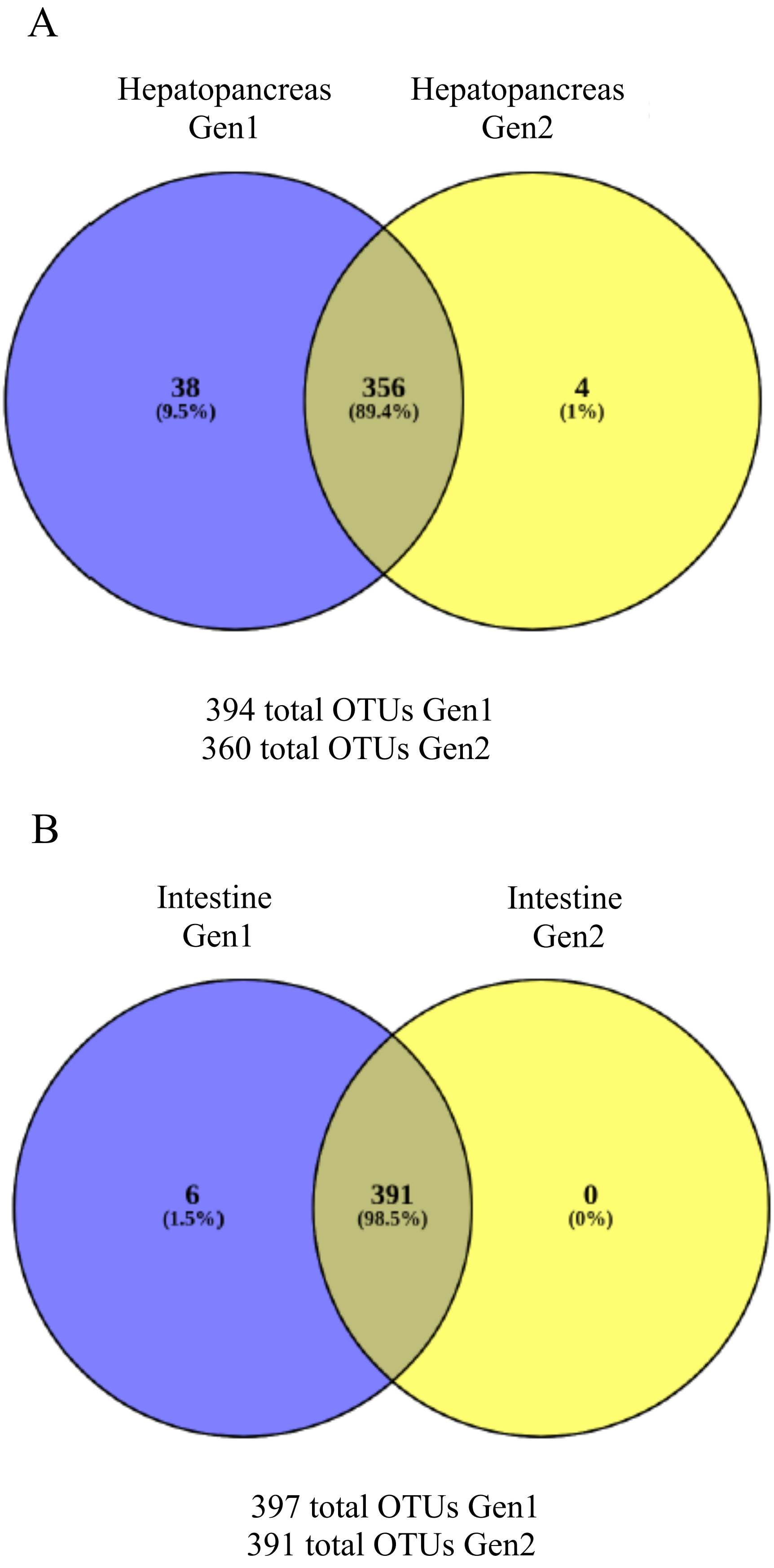


**Supplementary Figure 3. OTUs comparison between both genetic lines.** Venn diagrams comparing OTUs between both genetic lines in A) the hepatopancreas, and B) the intestine. The number of total OTUs is indicated below each diagram, and the corresponding percentage is indicated in parenthesis
